## Supplementary information for "Structure of a Rhs effector clade domain identifies new mechanistic insights into type VI secretion system toxin delivery"


| Item | Description | Pg |
|---|---|---|
| Fig S1 | Domain boundaries for Tse15. | 2 |
| Fig S2 | Workflow for solving Tse15 wild-type structure. | 3 |
| Fig S3 | Top AlphaFold2 model for Tse15. | 4 |
| Table S1 | Metrics for Tse15 structures | 5 |
| Fig S4 | Tse15 clade domain is common to many bacterial Rhs effectors. | 6 |
| Table S2 | List of homologues to the Tse15 N-terminal domain as determined by FoldSeek. | 7 |
| Fig S5 | Access to the interior of the Rhs cage and interactions with the clade domain. | 9 |
| Table S3 | Interactions between clade and Rhs domains in Tse15 | 10 |
| Fig S6 | Toxin difference maps for Tse15 and Tse15$_{NN}$ with fitted toxin residues. | 11 |
| Fig S7 | Density of toxin cleavage site in Tse15 and Tse15NN. | 12 |
| Fig S8 | Crosslinking mass spectrometry of Tse15. | 13 |
| Fig S9 | Tse15$_{NN}$ construct design and purification. | 14 |
| Fig S10 | Workflow for solving Tse15$_{NN}$ structure. | 15 |
| Fig S11 | Identification and mutation of clade autocleavage motif. | 16 |
| Table S4 | Conservation of the clade autocleavage motif. | 17 |
| Fig S12 | The VgrG15 N-terminal domain is highly conserved as compared to the N-terminal domains of the other *A. baumannii* AB307 VgrG proteins. | 18 |
| Fig S13 | Peptide signatures of Rhs effectors and their cognate VgrG within the secretome of T6SS active *A. baumannii.* | 19 |

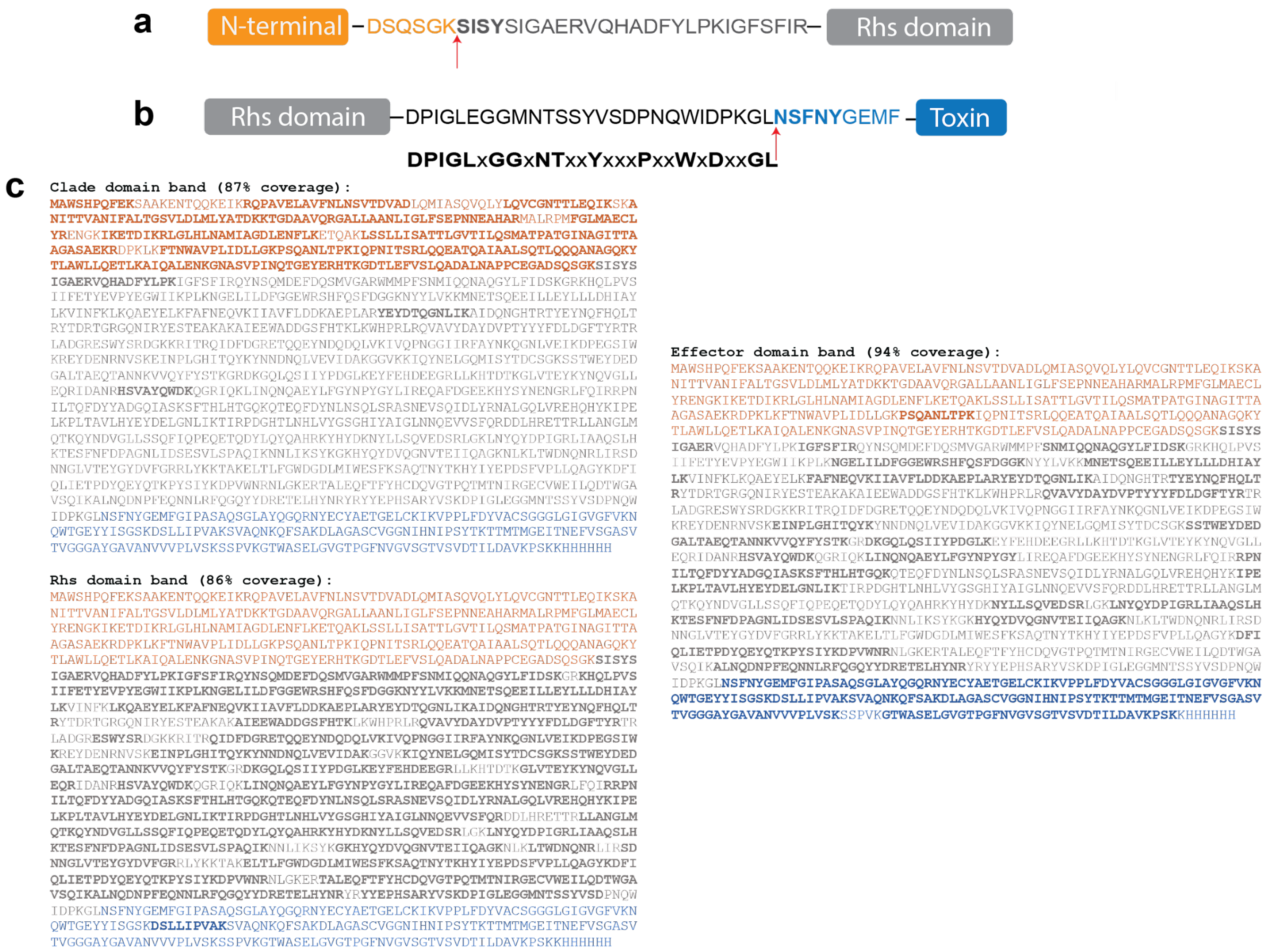


**Supp Figure 1. Domain boundaries for Tse15. a)** N-terminal sequencing of the Rhs domain. N-terminal domain is orange, Rhs domain is grey. Bold residues indicate residues identified by N-terminal sequencing. **b)** N-terminal sequencing of the toxin domain. Rhs domain is grey and toxin domain is blue. Blue bold residues indicate residues identified by N-terminal sequencing. Residues below show alignment of consensus sequence. **c)** Mass spectrometry peptide fingerprinting for excised bands of each domain from SDS-PAGE. Excised band is indicated at the top with percentage peptide coverage for each domain. Domains are coloured as follows: N-terminal clade residues are orange, Rhs domain residues grey and toxin domain residues blue. Residues shown in bold indicate peptides identified.

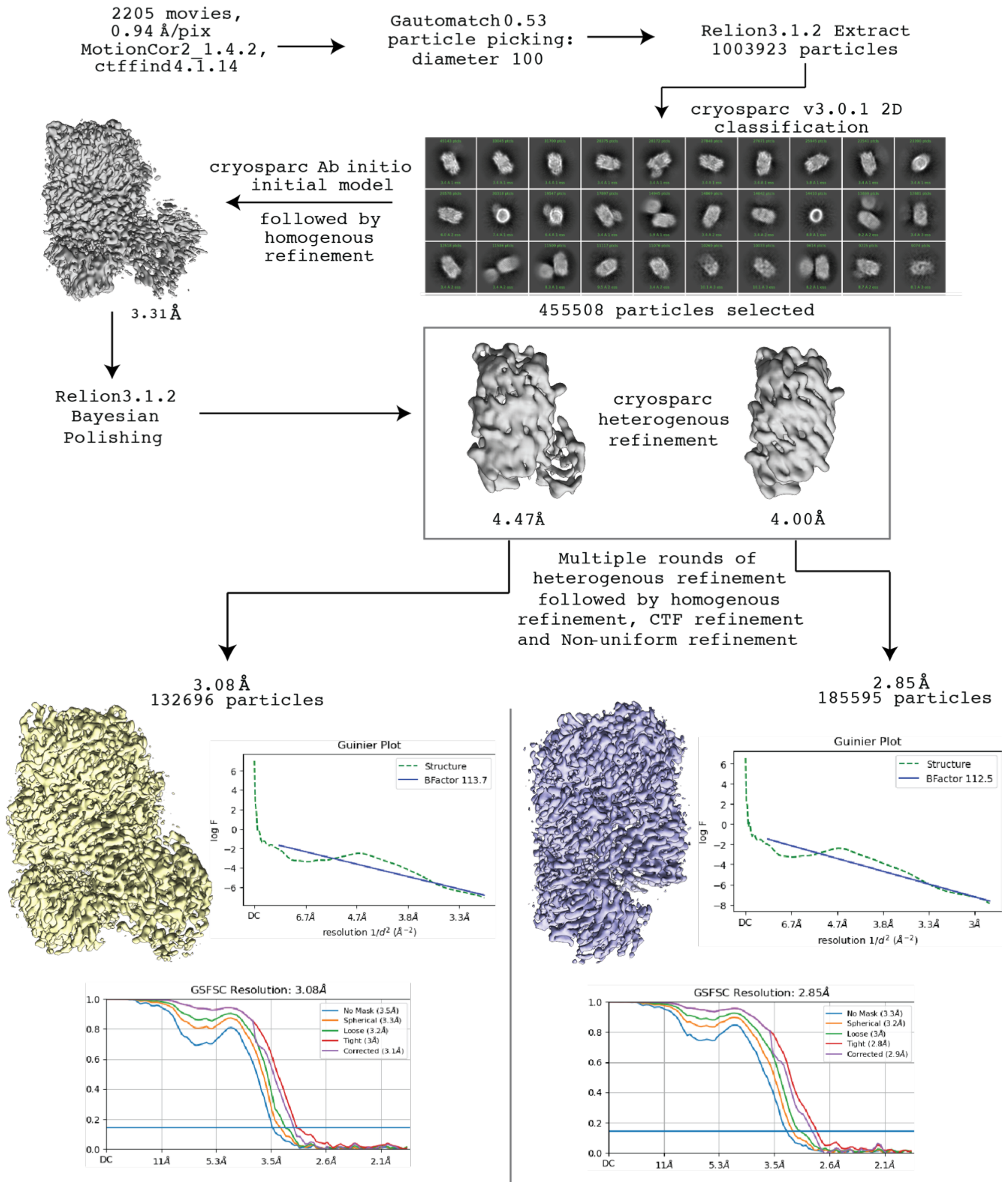


**Supp figure 2. Workflow for solving Tse15 wild-type structure.** Particles were picked using Gautomatch, then extracted using Relion. Cryosparc was used for 2D classification, and an *ab initio* model was produced, and homogenous refinement conducted producing a 3.31 Å map. Particles were polished in relion, and then used for heterogenous refinement in cryosparc to produce two particles classes at 4.47 and 4 Å. These were separately refined further using heterogenous refinement, CTF refinement and non-uniform refinement to produce a Tse15 map with the clade domain at 3.08 Å and a Tse15 map without the clade domain at 2.85 Å.

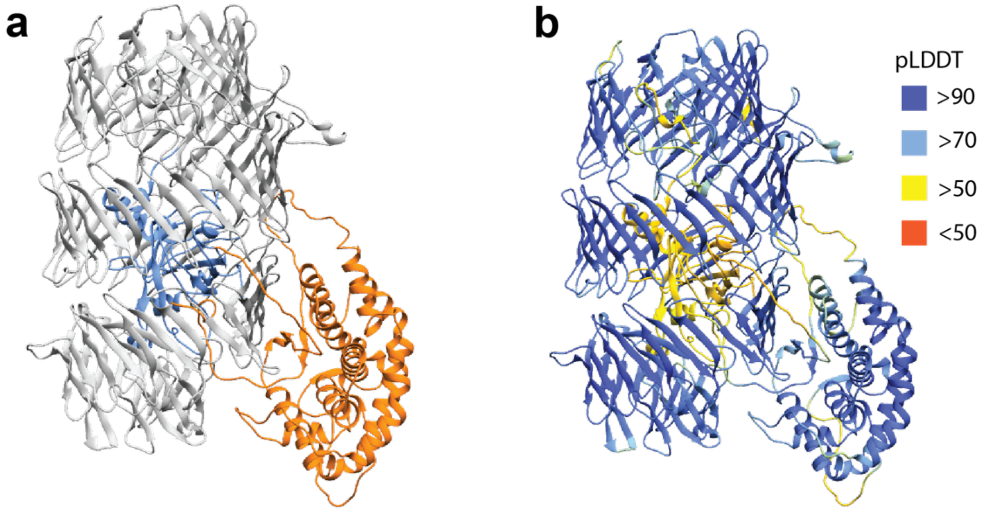


**Supp Figure 3. Top AlphaFold2 model for Tse15. a)** model with different domains coloured, orange is N-terminal domain, grey Rhs domain and blue toxin domain. **b)** pLDDT score mapped onto Tse15 model.

### Supp Table 1. Metrics for Tse15 structures.

| **Data collection** | **Tse15** | **Tse15$_{NN}$** |
|---|---|---|
| Particles | 132696 | 1,163,105 |
| Pixel size (Å) | 0.94 | 0.65 |
| Voltage (kV) | 200 | 300 |
| Electron dose (e/Å2) | 50 | 60 |
| **Refinement** | | |
| $CC_{map\_model}$ | 0.73 | 0.74 |
| **Model Quality** | | |
| RMSD Bond length (Å) | 0.03 | 0.003 |
| RMSD Bond angles (˚) | 0.756 | 0.480 |
| Ramachandran favoured (%) | 92.97 | 95.01 |
| Ramachandran outliers (%) | 0.42 | 0.50 |
| Ramachandran allowed (%) | 6.61 | 4.49 |
| Rotamer outliers (%) | 1.55 | 1.45 |
| C-beta deviations (%) | 2 | 0 |
| Clashscore | 7.71 | 4.58 |
| PDB / EMDB ID | 8UY4 / 42792 | 8XUT / 42775 |

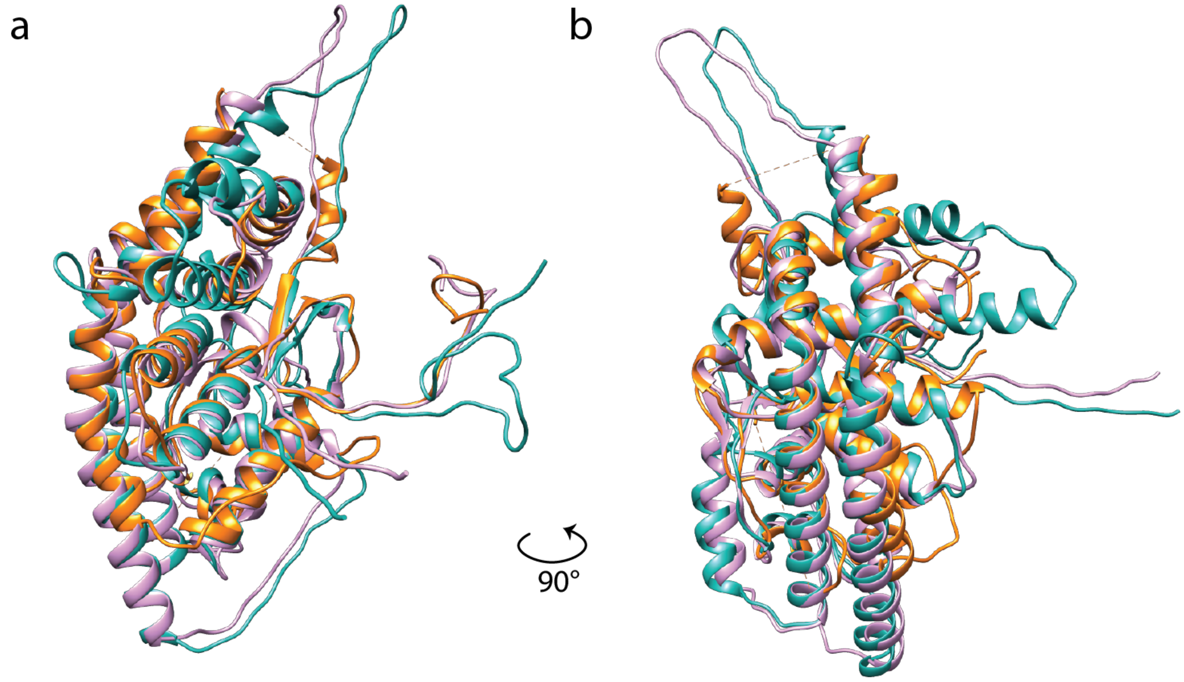


**Supp Figure 4. Tse15 clade domain is common to many bacterial Rhs effectors. a)** Cartoon alignment of the Tse15 N-terminal domain (orange) with a Rhs N-terminal domain from *Pseudomonas sp.* WS 5354 (pink, RMSD 7.2 Å: NCBI, WP_169966139.1) and *Burkholderia sp.* AU6039 (teal, RMSD 9.4 Å, NCBI: NKFK01000008.1).

**Supp Table 2. List of homologues to the Tse15 N-terminal domain as determined by FoldSeek[1].**

| Target | Description | Scientific Name | Prob. | Seq. Id. (%) | E-Value | Position in query |
|---|---|---|---|---|---|---|
| AF-A0A6I4HT81-F1-model_v4 | RHS repeat protein | *Acinetobacter baumannii* | 1 | 100 | 9.70E-46 | 1-333 |
| AF-L9LR64-F1-model_v4 | YD repeat protein (3 repeats) | *Acinetobacter sp.* OIFC021 | 1 | 94.5 | 1.61E-44 | 1-333 |
| AF-A0A7Z2HG68-F1-model_v4 | Rhs family protein | *Acinetobacter baumannii* | 1 | 98.1 | 9.45E-45 | 1-333 |
| AF-N9DJH2-F1-model_v4 | Uncharacterized protein | *Acinetobacter bereziniae* LMG 1003 = CIP 70.12 | 1 | 60.1 | 3.95E-34 | 1-326 |
| AF-A0A1H7XNA3-F1-model_v4 | YD repeat-containing protein | *Acinetobacter sp.* DSM 11652 | 1 | 58.7 | 2.48E-33 | 1-331 |
| AF-A0A429MRQ6-F1-model_v4 | RHS repeat protein | *Acinetobacter baumannii* | 1 | 100 | 6.89E-24 | 85-333 |
| AF-A0A7Y1FLN0-F1-model_v4 | RHS repeat protein | *Pseudomonas sp.* WS 5078 | 1 | 23.9 | 1.30E-14 | 1-332 |
| AF-A0A3M4WEW4-F1-model_v4 | RHS repeat containing protein | *Pseudomonas cichorii* | 1 | 25.9 | 3.42E-14 | 9-325 |
| AF-A0A519DSR3-F1-model_v4 | RHS repeat protein | *Pseudomonas sp.* | 1 | 25.5 | 4.80E-14 | 8-325 |
| **AF-A0A7Y1G8B5-F1-model_v4** | **RHS repeat protein** | ***Pseudomonas sp.* WS 5354** | **1** | **24.7** | **6.41E-14** | **2-325** |
| AF-A0A1I0LVF7-F1-model_v4 | YD repeat-containing protein | *Burkholderia cepacia* | 1 | 24.3 | 4.80E-14 | 1-311 |
| AF-A0A4R7US80-F1-model_v4 | Uncharacterized protein | *Pseudomonas helmanticensis* | 1 | 25.3 | 3.59E-14 | 5-325 |
| AF-A0A7C1WQ62-F1-model_v4 | RHS repeat protein | *Pseudomonas graminis* | 1 | 22.8 | 3.49E-13 | 1-333 |
| AF-A0A3P0RKZ5-F1-model_v4 | RHS repeat protein | *Burkholderia sp.* Bp9004 | 1 | 23.4 | 3.17E-13 | 1-311 |
| AF-A0A519FHV8-F1-model_v4 | RHS repeat protein | *Pseudomonas sp.* | 1 | 23.7 | 1.73E-12 | 12-322 |
| AF-A0A107A4U9-F1-model_v4 | Uncharacterized protein | *Burkholderia stagnalis* | 1 | 22.8 | 1.35E-12 | 7-325 |
| AF-J2TDQ0-F1-model_v4 | Rhs family protein | *Variovorax sp.* CF313 | 1 | 23.6 | 3.75E-12 | 1-327 |
| AF-A0A365QG90-F1-model_v4 | RHS family protein | *Burkholderia reimsis* | 1 | 21.5 | 3.75E-12 | 15-333 |
| AF-A0A7X2BG59-F1-model_v4 | RHS repeat protein | *Pseudomonas sp.* FSL R10-0765 | 1 | 22.7 | 9.40E-12 | 12-332 |
| AF-A0A837MJY1-F1-model_v4 | Uncharacterized protein | *Pseudomonas helleri* | 1 | 24 | 6.85E-11 | 12-324 |
| AF-A0A6I4HSU2-F1-model_v4 | Type IV secretion protein Rhs | *Acinetobacter baumannii* | 1 | 22.5 | 2.04E-11 | 1-322 |
| AF-A0A7Y1HMJ9-F1-model_v4 | RHS repeat protein | *Pseudomonas lundensis* | 1 | 23.8 | 5.12E-11 | 21-326 |
| AF-A0A1H7LLH1-F1-model_v4 | Uncharacterized protein | *Variovorax sp.* YR750 | 1 | 24.9 | 2.36E-11 | 8-333 |
| AF-A0A7G2MP46-F1-model_v4 | Uncharacterized protein | *Pseudomonas sp.* | 1 | 23.3 | 2.19E-10 | 12-324 |
| AF-A0A4R8LT33-F1-model_v4 | YD repeat-containing protein | *Paraburkholderia rhizosphaerae* | 1 | 22.1 | 1.08E-09 | 5-333 |
| AF-A0A1H7ETW9-F1-model_v4 | Uncharacterized protein | *Variovorax sp.* OK202 | 1 | 25.2 | 3.65E-11 | 1-333 |
| AF-A0A6L5HR91-F1-model_v4 | Uncharacterized protein | *Pseudomonas helleri* | 1 | 23.3 | 2.30E-10 | 10-318 |
| AF-A0A3P0RGT4-F1-model_v4 | Uncharacterized protein | *Burkholderia sp.* Bp9004 | 1 | 23.5 | 3.91E-10 | 31-333 |
| AF-A0A4Y8HF87-F1-model_v4 | YD repeat-containing protein | *Pseudomonas sp.* URIL14HWK12:I1 | 1 | 20.8 | 2.59E-09 | 7-315 |
| AF-A0A1W1Y1F1-F1-model_v4 | YD repeat-containing protein | *Andreprevotia lacus* DSM 23236 | 1 | 22.8 | 2.59E-09 | 8-331 |
| AF-A0A071MKM1-F1-model_v4 | Sugar-binding protein | *Burkholderia cenocepacia* | 1 | 19.9 | 1.41E-08 | 17-321 |
| **AF-A0A228NXD1-F1-model_v4** | **Type IV secretion protein Rhs** | ***Burkholderia sp.* AU6039** | **1** | **19.8** | **1.45E-09** | **7-333** |

| | | | | | | |
|---|---|---|---|---|---|---|
| AF-N9MJ93-F1-model_v4 | Uncharacterized protein | *Acinetobacter sp.* ANC 4105 | 1 | 83.4 | 3.34E-15 | 115-332 |
| AF-A0A358GY28-F1-model_v4 | Type IV secretion protein Rhs | *Acinetobacter sp.* | 1 | 20.2 | 4.29E-08 | 17-327 |
| AF-A0A3P0RK35-F1-model_v4 | RHS repeat protein | *Burkholderia sp.* Bp9004 | 1 | 17.5 | 1.83E-07 | 17-311 |
| AF-N8WS81-F1-model_v4 | Uncharacterized protein | *Acinetobacter guillouiae* NIPH 991 | 1 | 18.9 | 2.34E-07 | 12-326 |
| AF-A0A8B5S1J0-F1-model_v4 | RHS repeat protein | *Acinetobacter bereziniae* | 1 | 60.2 | 7.03E-12 | 106-333 |
| AF-A0A4R6Y000-F1-model_v4 | YD repeat-containing protein | *Hydromonas duriensis* | 1 | 17.8 | 3.13E-07 | 1-324 |
| AF-A0A009GBA8-F1-model_v4 | RHS Repeat family protein | *Acinetobacter baumannii* 118362 | 1 | 14.9 | 3.79E-07 | 17-320 |
| AF-A0A429MUJ9-F1-model_v4 | RHS repeat protein | *Acinetobacter baumannii* | 1 | 15.4 | 6.78E-07 | 17-320 |
| AF-A0A4R3W625-F1-model_v4 | YD repeat-containing protein | *Pseudomonas sp.* LP_8_YM | 1 | 17 | 9.27E-06 | 16-315 |
| AF-A0A3N7ZDS0-F1-model_v4 | RHS repeat protein | *Burkholderia sp.* Bp9142 | 1 | 19.4 | 1.07E-05 | 7-328 |
| AF-A0A7X1WP70-F1-model_v4 | Rhs family protein | *Pseudomonas sp.* FSL R10-0399 | 1 | 18.3 | 2.28E-06 | 8-324 |
| AF-A0A6P2ZV05-F1-model_v4 | Sugar-binding protein | *Burkholderia contaminans* | 1 | 17.8 | 3.11E-05 | 27-325 |
| AF-L9MNM7-F1-model_v4 | Uncharacterized protein | *Acinetobacter sp.* WC-743 | 1 | 61.7 | 1.97E-06 | 1-159 |
| AF-A0A3S0B389-F1-model_v4 | RHS repeat protein | *Variovorax sp.* 679 | 1 | 22.3 | 5.84E-05 | 94-302 |
| AF-A0A3P0RI29-F1-model_v4 | RHS repeat protein | *Burkholderia sp.* Bp9004 | 1 | 18.9 | 1.50E-05 | 7-299 |
| AF-A0A1H7QHI4-F1-model_v4 | Uncharacterized protein | *Variovorax sp.* YR750 | 1 | 23.9 | 7.09E-05 | 1-299 |
| AF-A0A7L7VH69-F1-model_v4 | Uncharacterized protein | *Pseudomonas putida* | 1 | 16.1 | 8.79E-04 | 17-326 |
| AF-A0A0D0MNU1-F1-model_v4 | Uncharacterized protein | *Variovorax paradoxus* | 1 | 20.4 | 7.60E-04 | 21-290 |
| AF-A0A6N8T242-F1-model_v4 | Sugar-binding protein | *Burkholderia sp.* 4701 | 1 | 13.9 | 1.47E-04 | 25-302 |
| AF-J3CCY2-F1-model_v4 | Rhs family protein | *Variovorax sp.* CF313 | 1 | 23 | 1.02E-03 | 99-333 |
| AF-A0A7X1W4E1-F1-model_v4 | Rhs family protein | *Pseudomonas sp.* FSL R10-0765 | 1 | 17.4 | 1.50E-03 | 26-333 |
| AF-A0A2U9SMH5-F1-model_v4 | Uncharacterized protein | *Burkholderia sp.* JP2-270 | 1 | 16.1 | 9.23E-04 | 21-326 |
| AF-A0A4R0X6U5-F1-model_v4 | Uncharacterized protein | *Paraburkholderia steynii* | 1 | 16.5 | 1.43E-03 | 13-333 |
| AF-A0A105TLI1-F1-model_v4 | Uncharacterized protein | *Pseudomonas sp.* TAD18 | 0.93 | 14.2 | 8.33E-02 | 26-333 |
| AF-A0A143IY85-F1-model_v4 | Uncharacterized protein | *Acinetobacter pittii* | 0.72 | 14.6 | 2.20E-01 | 21-333 |
| AF-S9QIT6-F1-model_v4 | Filamentous hemagglutinin | *Cystobacter fuscus* DSM 2262 | 0.6 | 11.5 | 6.37E-01 | 17-333 |
| AF-A0A3N8B1N8-F1-model_v4 | RHS family protein | *Burkholderia sp.* Bp9143 | 0.57 | 19.4 | 1.32E+00 | 152-332 |
| AF-E0SGL7-F1-model_v4 | Putative deoxyribonuclease RhsC | *Dickeya dadantii* 3937 | 0.54 | 11.4 | 5.78E-01 | 68-333 |
| AF-A0A6I3T6V6-F1-model_v4 | Type IV secretion protein Rhs | *Pseudoduganella buxea* | 0.44 | 19 | 1.38E+00 | 146-333 |
| AF-N9RKC7-F1-model_v4 | Uncharacterized protein | *Acinetobacter sp.* NIPH 2100 | 0.35 | 8.6 | 1.52E+00 | 34-333 |
| AF-A0A7H2WCG2-F1-model_v4 | Ntox15 domain-containing protein | *Acinetobacter seifertii* | 0.3 | 12.9 | 2.04E+00 | 26-333 |
| AF-A0A378QIP5-F1-model_v4 | Uncharacterized protein | *Moraxella lacunata* | 0.23 | 11.3 | 4.64E+0 | 30-333 |

**Bold** indicates structures used in alignment in **Supp Figure 4**.

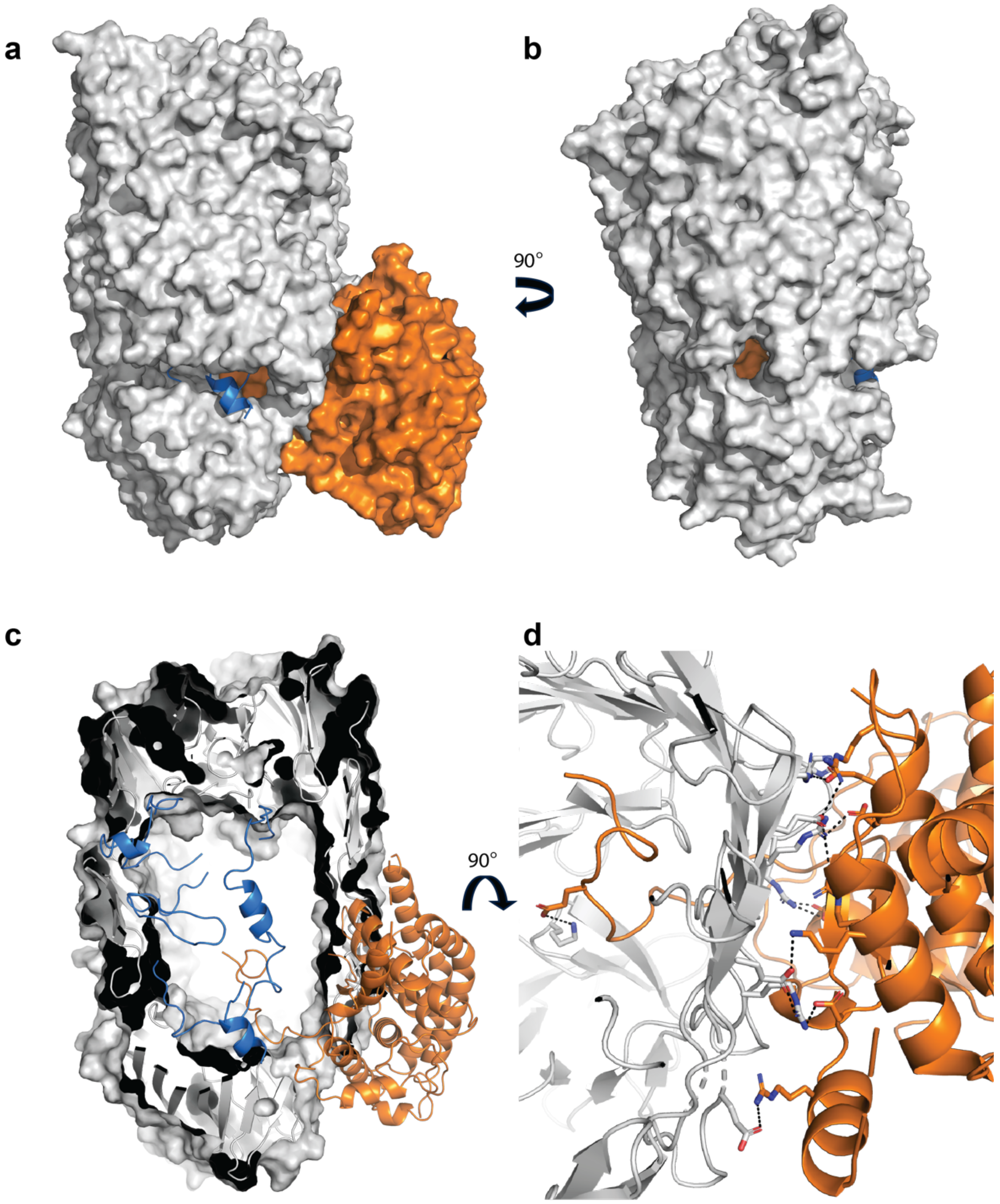


**Supp Figure 5. Access to the interior of the Rhs cage and interactions with the clade domain.** Tse15 is shown in surface fill with the Rhs domain in grey and clade in orange. The toxin is shown in cartoon with carbon atoms colored blue. **a**) and **b**) Access to the interior of the cage occurs via an opening between the bottom and middle b-stranded substructures. The toxin can be seen in the opening. **c**) Through the opening, the clade domain can be visualised to be well-coordinated with the Rhs domain, as well as close to the toxin density. **d**) Salt bridges dominate the protein-protein interface of the Rhs:clade interaction where both are shown in cartoon, with Rhs carbon atoms in grey and clade in orange. Residues forming salt-bridges are shown in stick.

**Supplementary Table 3: Interactions between clade and Rhs domains in Tse15.** Generated by PDBePISA[2].

| | Clade residue [atom] | Distance (Å) | Rhs residue [atom] |
|---|---|---|---|
| **Salt bridges** | | | |
| 1 | GLU18 [OE1] | 2.81 | HIS589 [NE2] |
| 2 | GLU18 [OE2] | 2.93 | ARG619 [NH2] |
| 3 | ASP32 [OD1] | 2.78 | ARG593 [NH2] |
| 4 | ASP32 [OD1] | 2.90 | ARG593 [NH1] |
| 5 | ASP32 [OD2] | 3.04 | ARG593 [NH2] |
| 6 | GLU127 [OE1] | 2.91 | ARG617 [NH1] |
| 7 | GLU127 [OE1] | 3.30 | ARG617 [NH2] |
| 8 | ASP308 [OD2] | 2.49 | ARG638 [NE] |
| 9 | ASP308 [OD2] | 2.84 | ARG638 [NH2] |
| 10 | GLU326 [OE2] | 3.94 | LYS400 [NZ] |
| 11 | ARG131 [NH1] | 3.47 | GLU625 [OE2] |
| 12 | LYS142 [NZ] | 3.41 | GLU646 [OE2] |
| 13 | ARG143 [NH2] | 3.76 | GLU625 [OE1] |
| 14 | ARG303 [NE] | 2.90 | GLU650 [OE1] |
| 15 | ARG303 [NH2] | 3.90 | GLU650 [OE1] |
| **Hydrogen bonds** | | | |
| 1 | GLU18 [OE2] | 1.78 | TYR608 [HH] |
| 2 | GLU18 [OE2] | 2.04 | GLN594 [HE22] |
| 3 | GLU18 [OE2] | 2.45 | ARG619 [HH22] |
| 4 | ASP32 [OD1] | 2.33 | ARG593 [HH22] |
| 5 | ASP32 [OD1] | 2.47 | ARG593 [HH12] |
| 6 | GLU127 [OE1] | 2.12 | ARG617 [HH12] |
| 7 | GLU138 [OE2] | 2.47 | ASN889 [HD21] |
| 8 | ASP308 [OD2] | 1.83 | ARG638 [HE] |
| 9 | ASP308 [OD2] | 2.26 | ARG638 [HH21] |
| 10 | ASP308 [OD2] | 3.89 | SER629 [OG] |
| 11 | GLN316 [OE1] | 2.48 | ARG593 [H] |
| 12 | ALA322 [O] | 1.93 | ARG402 [HE] |
| 13 | ALA322 [O] | 2.41 | ARG402 [HH21] |
| 14 | LYS137 [HZ1] | 1.63 | PHE642[ O] |
| 15 | LYS142 [HZ1] | 2.29 | GLY644 [O] |
| 16 | ARG303 [HE] | 2.26 | GLU650 [OE1] |
| 17 | GLN316 [H] | 2.22 | ARG591 [O] |
| 18 | LEU320 [H] | 2.02 | LEU611 [O] |
| 19 | SER332[H] | 2.23 | ILE336[O] |

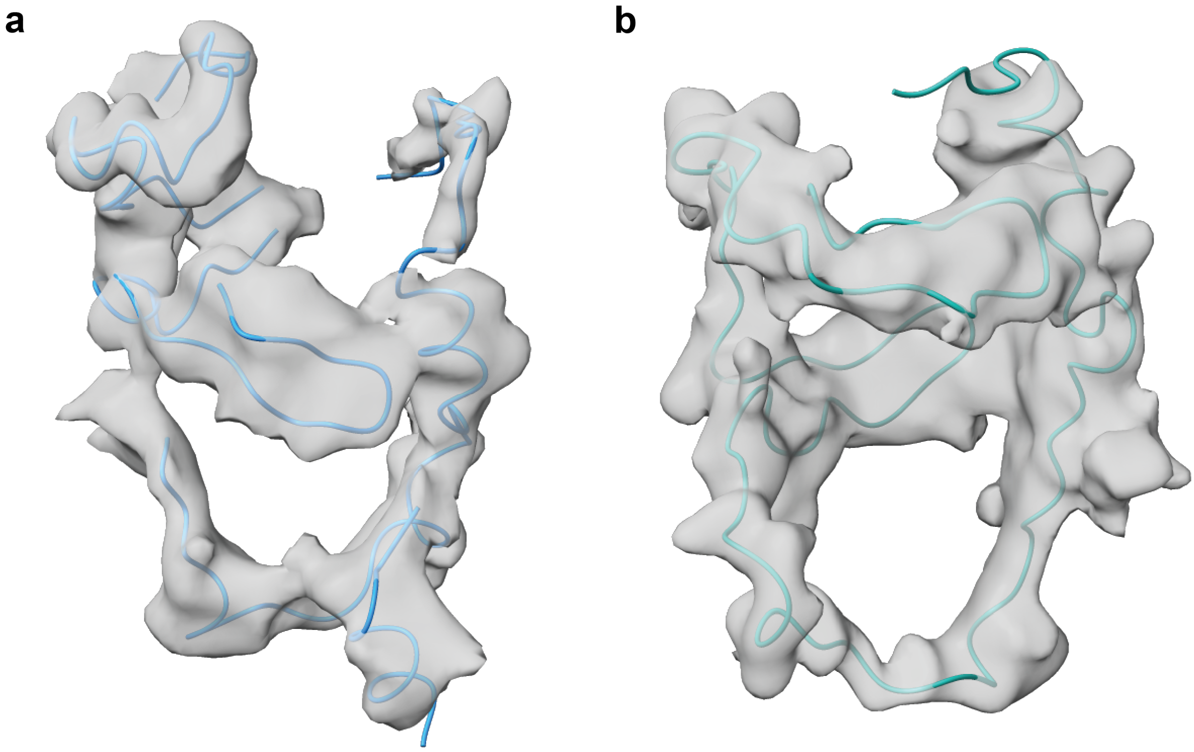


**Supp Figure 6. Toxin difference maps for Tse15 and Tse15$_{NN}$ with fitted toxin residues.** Toxin density for both maps is shown in grey, and is set to show density around the chains at a range of 2.5 on Chimera X. **a)** Tse15 with the toxin shown in blue, map set at an RMSD of 9.7 Å and b) Tse15$_{NN}$ with the toxin shown in cyan, map set at an RMSD of 5.1 Å.

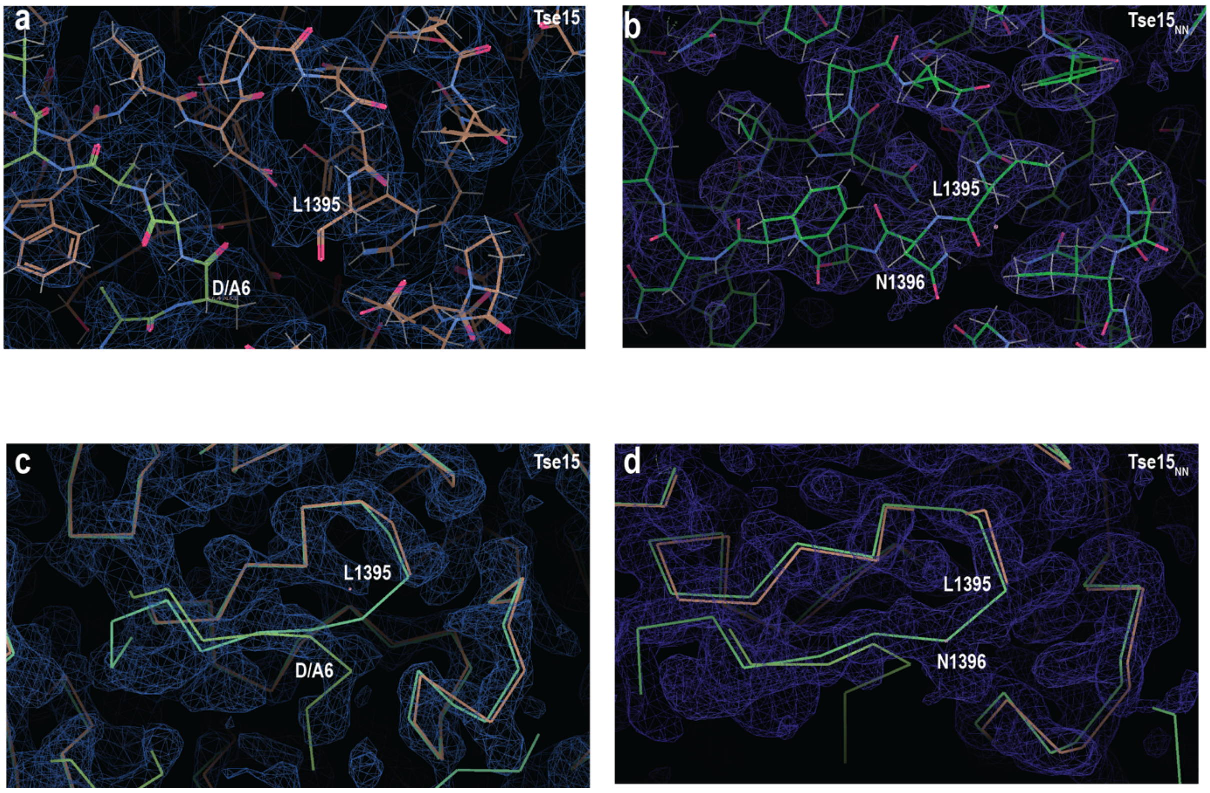


**Supp Figure 7. Density of toxin cleavage site in Tse15 and Tse15$_{NN}$.** Carbon atoms show Tse15 in bone (chain A) and green (chain D); Tse15$_{NN}$ (chain A) in dark green. Coulomb potential maps for Tse15 (**a, c**; r.m.s.d. contour 6.0) and Tse15$_{NN}$ (**b, d**; r.m.s.d. contour 5.0) are shown in blue. The position of toxin autocleavage, L1395 is indicated as is the next residue N1396 for Tse15$_{NN}$. The position of toxin peptide from the D-chain of Tse15 is indicated as A6.

**a**

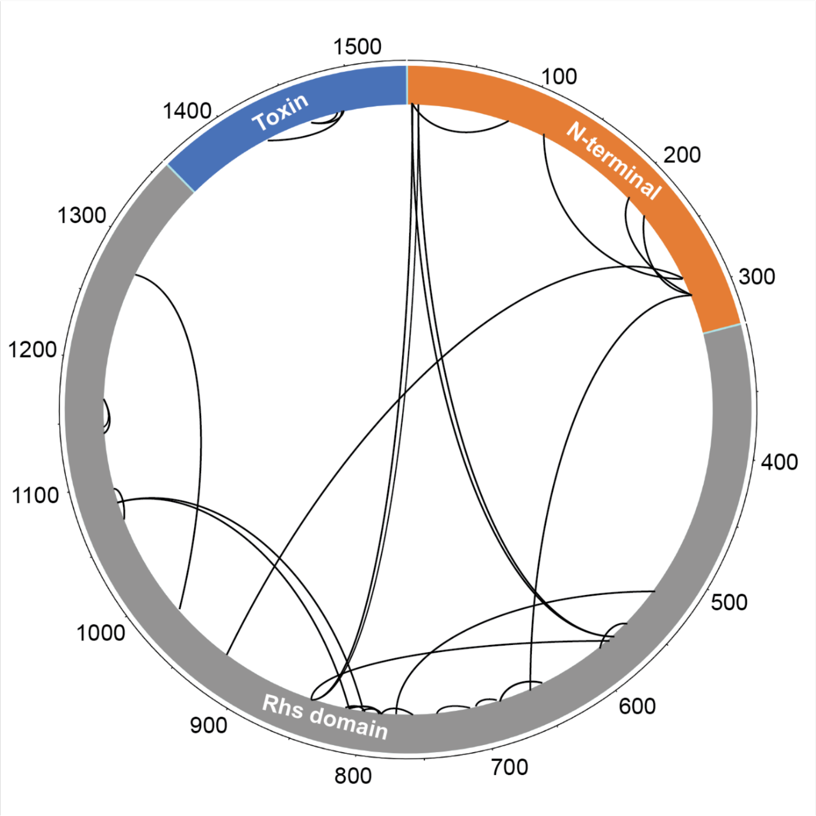


**b**

| Crosslinked residues | |
|---|---|
| 3 | 846 |
| 3 | 587 |
| 9 | 85 |
| 9 | 587 |
| 9 | 846 |
| 137 | 288 |
| 210 | 306 |
| 225 | 306 |
| 288 | 925 |
| 306 | 659 |
| 512 | 784 |
| 569 | 587 |
| 571 | 587 |
| 585 | 846 |
| 659 | 682 |
| 690 | 701 |
| 713 | 732 |
| 771 | 784 |
| 780 | 797 |
| 780 | 810 |
| 797 | 810 |
| 797 | 1077 |
| 810 | 784/797 |
| 815 | 1077 |
| 971 | 1279 |
| 1072 | 1091 |
| 1140 | 1166 |
| 1145 | 1166 |
| 1434 | 1491 |
| 1471 | 1486 |
| 1471 | 1491 |

**Supp Figure 8. Crosslinking mass spectrometry of Tse15.** For both a and b, orange indicates N-terminal domain, grey Rhs domain and blue toxin domain. **a)** crosslinks mapped onto the Tse15 sequence. Residue numbers are indicated on the outside, domains are coloured and labelled. **b)** crosslinks in list form for clarity.

**a**

```
Tse15    HSARYVSKDPIGLEGGMNTSSYVSDPNQWIDPKGLNSFNY 1400
Tse15NN  HSARYVSKNPIGLEGGMNTSSYVSDPNQWINPKGLNSFNY 1400
         ********:********************:*********
```

**b**

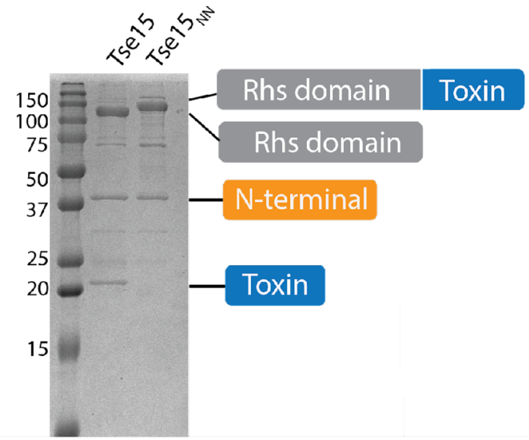


**c**

**Tse15NN Rhs and toxin domain band (92% coverage):**

```
MAWSHPQFEKSAAKENTQQKEIKRQPAVELAVFNLNSVTDVADLQMIASQVQLYLQVCGNTTLEQIKSKANITTVANIFALTGSVLDLMLYATDKKTGDAAVQRG
ALLAANLIGLFSEPNNEAHARMALRPMFGLMAECLYRENGKIKETDIKRLGLHLNAMIAGDLENFLKETQAKLSSLLISATTLGVTILQSMATPATGINAGITTA
AGASAEKRDPKLKFTNWAVPLIDLLGKPSQANLTPKIQPNITSRLQQEATQAIAALSQTLQQQANAGQKYTLAWLLQETLKAIQALENKGNASVPINQTGEYERH
TKGDTLEFVSLQADALNAPPCEGADSQSGKSISYSIGAERVQHADFYLPKIGFSFIRQYNSQMDEFDQSMVGARWMMPFSNMIQQNAQGYLFIDSKGRKHQLPVS
IIFETYEVPYEGWIIKPLKNGELILDFGGEWRSHFQSFDGGKNYYLVKKMNETSQEEILLEYLLLDHIAYLKVINFKLKQAEYELKFAFNEQVKIIAVFLDDKAE
PLARYEYDTQGNLIKAIDQNGHTRTYEYNQFHQLTRYTDRTGRGQNIRYESTEAKAKAIEEWADDGSFHTKLKWHPRLRQVAVYDAYDVPTYYYFDLDGFTYRTR
LADGRESWYSRDGKKRITRQIDFDGRETQQEYNDQDQLVKIVQPNGGIIRFAYNKQGNLVEIKDPEGSIWKREYDENRNVSKEINPLGHITQYKYNNDNQLVEVI
DAKGGVKKIQYNELGQMISYTDCSGKSSTWEYDEDGALTAEQTANNKVVQYFYSTKGRDKGQLQSIIYPDGLKEYFEHDEEGRLLKHTDTKGLVTEYKYNQVGLL
EQRIDANRHSVAYQWDKQGRIQKLINQNQAEYLFGYNPYGYLIREQAFDGEEKHYSYNENGRLFQIRRPNILTQFDYYADGQIASKSFTHLHTGQKQTEQFDYNL
NSQLSRASNEVSQIDLYRNALGQLVREHQHYKIPELKPLTAVLHYEYDELGNLIKTIRPDGHTLNHLVYGSGHIYAIGLNNQEVVSFQRDDLHRETTRLLANGLM
QTKQYNDVGLLSSQFIQPEQETQDYLQYQAHRKYHYDKNYLLSQVEDSRLGKLNYQYDPIGRLIAAQSLHKTESFNFDPAGNLIDSESVLSPAQIKNNLIKSYKG
KHYQYDVQGNVTEIIQAGKNLKLTWDNQNRLIRSDNNGLVTEYGYDVFGRRLYKKTAKELTLFGWDGDLMIWESFKSAQTNYTKHYIYEPDSFVPLLQAGYKDFI
QLIETPDYQEYQTKPYSIYKDPVWNRNLGKERTALEQFTFYHCDQVGTPQTMTNIRGECVWEILQDTWGAVSQIKALNQDNPFEQNNLRFQGQYYDRETELHYNR
YRYYEPHSARYVSKNPIGLEGGMNTSSYVSDPNQWINPKGLNSFNYGEMFGIPASAQSGLAYQGQRNYECYAETGELCKIKVPPLFDYVACSGGGLGIGVGFVKN
QWTGEYYISGSKDSLLIPVAKSVAQNKQFSAKDLAGASCVGGNIHNIPSYTKTTMTMGEITNEFVSGASVTVGGGAYGAVANVVVPLVSKSSPVKGTWASELGVG
TPGFNVGVSGTVSVDTILDAVKPSKKHHHHHH
```

**Supp Figure 9. Tse15NN construct design and purification. a)** sequence alignment showing mutation site for Tse15NN. Mutation sites are indicated in red, while blue indicates the start of the toxin domain. **b)** Coomassie stained SDS-PAGE gel showing purified Tse15NN compared to Tse15 wild-type. Domains as determined by mass spectrometry are shown. **c)** Mass spectrometry peptide fingerprinting analysis of the Tse15NN Rhs and toxin domain band excised from SDS-PAGE. Protein domains are coloured as follows: N-terminal clade residues orange, Rhs domain residues grey and toxin residues blue. Percentage peptide coverage is indicated at top. Residues shown in bold indicate peptides identified.

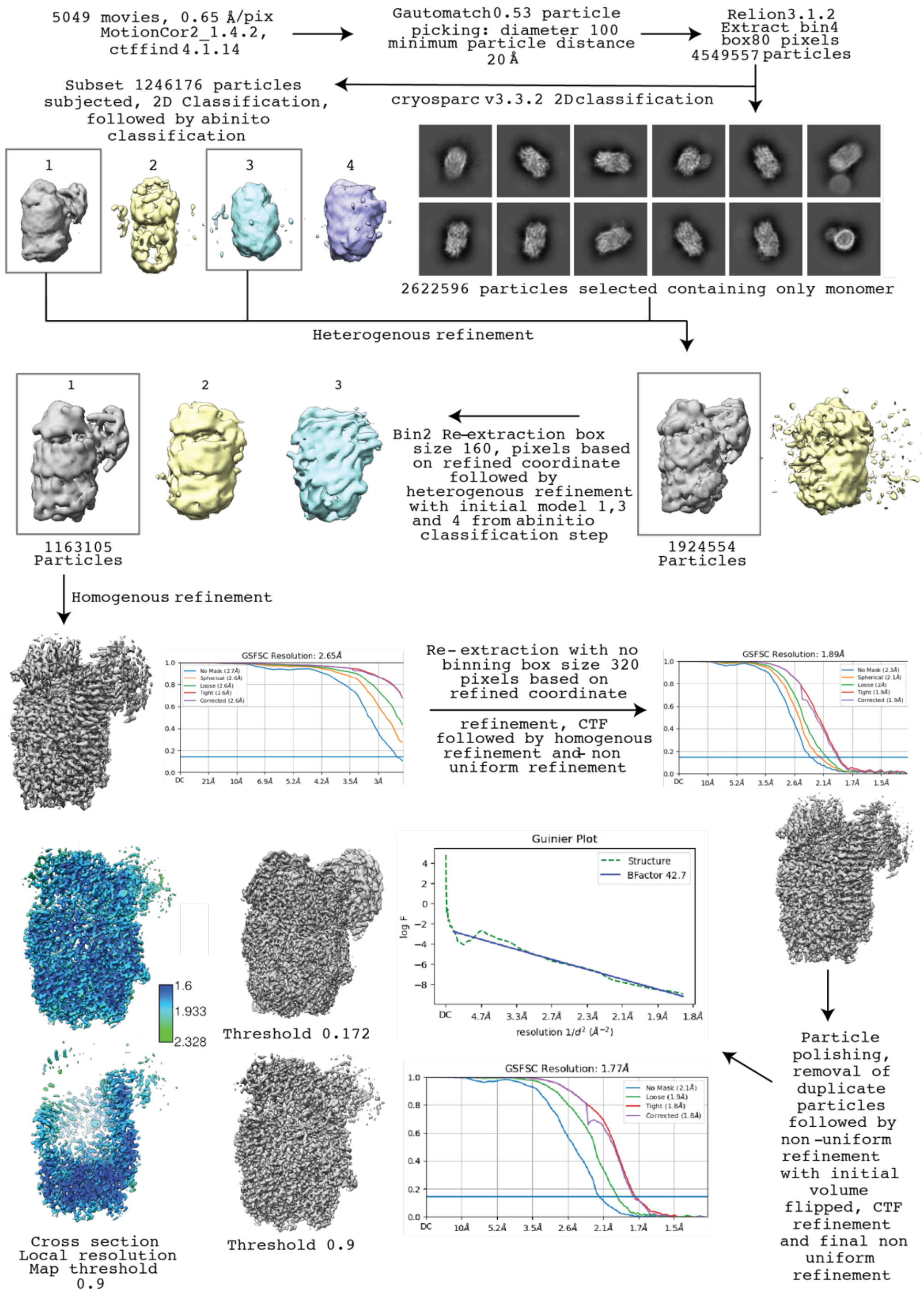


**Supp figure 10. Workflow for solving Tse15$_{NN}$ structure.** Particles were picked using Gautomatch, then extracted using Relion. Cryosparc was used for 2D classification, and four different *ab initio* models were produced. Heterogenous refinement was conducted using *ab initio* maps 1 and 3. Particles were re-extracted using a large box size, and heterogenous refinement was conducted using initial models 1, 3 and 4 from the *ab initio* reconstruction. Model 1 was then refined using homogenous refinement to produce a map at 2.65 Å, particles were then re-extracted based on refined coordinate refinement, CTF refinement, followed by both heterogenous refinement and non-uniform refinement to produce a map domain at 1.89 Å. Particles were polished, duplicate particles moved and then non-uniform refinement was conducted alongside CTF refinement and a last non-uniform refinement to produce a final model at 1.77 Å. Final maps shown at a threshold of 0.172 and 0.9. Local resolution is also shown at a map threshold of 0.9.

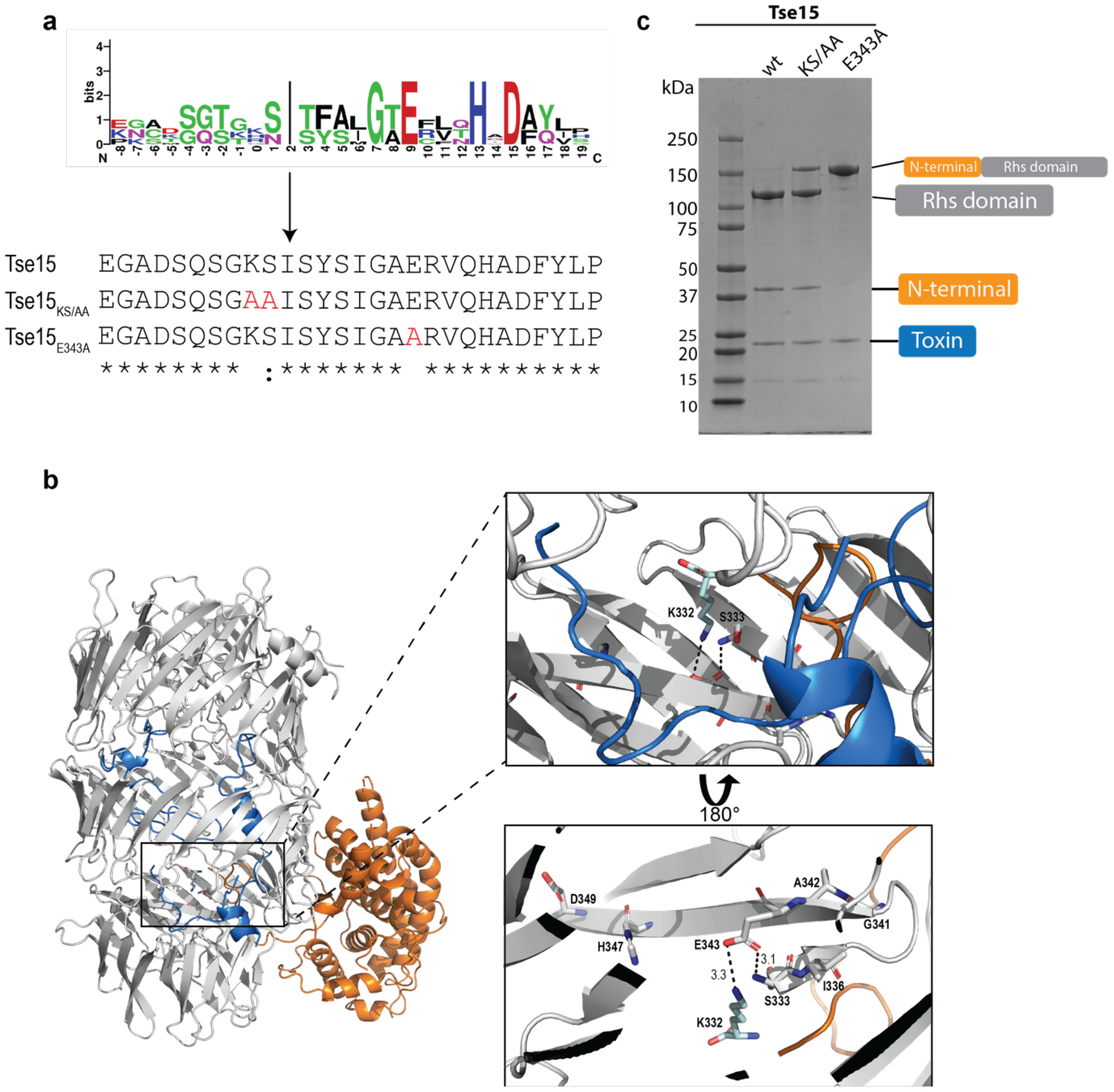


**Supp Figure 11. Identification and mutation of clade autocleavage motif. a)** *Acinetobacter spp* N-terminal cleavage site consensus shown as a sequence logo. Residues are coloured by property. Below the consensus is an alignment of the sequence of cleavage site in Tse15 structure and positions of KS and E343A mutations introduced. **b**) Location of the cleavage motif shown on the structure of Tse15 colored by domain, clade orange, Rhs grey and toxin blue. The inset zoom box shows the motif residues shown as stick with K332 provided by the Alphafold2 model and shown in cyan carbon stick. Below box is the same structure rotated 180° to show position of motif residues and interactions of nucleophilic E343. **c**) Coomassie stained SDS-PAGE showing N-terminal cleavage mutations, boxes to the right indicate domains.

**Supp Table 4. Conservation of the clade autocleavage motif across N-terminal homologues.** Searching FoldSeek[1] for homologous sequences identified that the autocleavage motif is highly conserved. The motif extent within Tse15 is shown in bold with conserved motif residues shown in red. Sequences above the black line had a probability score of $\geq 0.84$. Sequences below the line did not show the motif but had probability scores of $\leq 0.35$.

| | | |
|---|---|---|
| **Tse15** | **GKSISYSIGAERVQHADFY-LP-KIGFSFIRQYN----SQMDEFD-QSMV----------** | **375** |
| *Paraburkholderia_steynii* | GGSINYVMGDENLEQTDFA-LDGVVPIVWTRLYR----SSLVAYD-GSVL---------- | 407 |
| *Acinetobacter_bereziniae*_LMG_1003 | EHSISYAIGAERVNHADFY-LP-KLGFAFSRQYN----SQMNEFD-YSMI---------- | 388 |
| *Acinetobacter*_sp._WC-743 | ------------------------------------------------------------ | 172 |
| *Acinetobacter*_sp._DSM_11652 | AHSISYSIGAERVSHVDFS-LP-KLGLVFNRLYN----SQMAEFD-NGAL---------- | 379 |
| *Acinetobacter*_sp._OIFC021 | GKSISYSIGAEWVQHADFY-LP-KIGFSFIRQYN----SQMDEFD-QSMV---------- | 375 |
| *Acinetobacter_baumannii*_b | GKSISYSIGAERVQHADFY-LP-KIGFSFTRQYN----SQMDEFD-QSMV---------- | 375 |
| *Acinetobacter_baumannii*_c | GKSISYSIGAERVQHADFY-LP-KIGFSFIRQYN----SQMDEFD-QSMV---------- | 290 |
| *Acinetobacter_baumannii*_a | GKSISYSIGAERVQHADFY-LP-KIGFSFIRQYN----SQMDEFD-QSMV---------- | 375 |
| *Variovorax_paradoxus* | ------------------------------------------------------------ | 328 |
| *Hydromonas_duriensis* | TCDINFAMGVERVFHEDFS-IGHVLPIEWVRTYS----SAYAQYD-QSVM---------- | 444 |
| *Variovorax*_sp._CF313_a | GGAISLATGCESFGHTDFV-LGAPLPITWTRTYR----SDLAAFD-QGSL---------- | 393 |
| *Variovorax*_sp._679 | GGSISLVTGCESFTHTDFV-LEAPLPIEWARTYR----SQLDAYD-RGVL---------- | 298 |
| *Variovorax*_sp._CF313_b | GASISLSTGNESFTHTDFV-LAAPLPIEWARTYA----SDLDAFD-RSSL---------- | 293 |
| *Variovorax*_sp._YR750_a | RFAISLATGCESFTHVDFA-LVAPMPITWARTYR----SNLGAFD-EGSL---------- | 414 |
| *Variovorax*_sp._OK202 | GASISLATGNESFMHTDFV-LA-------------------------------------- | 375 |
| *Variovorax*_sp._YR750_b | ------------------------------------------------------------ | 315 |
| *Pseudomonas*_sp._VLB120 | GRTVHYLYEPGSFVPVAQALRKRPIVLHRQPDWR----QREYDFD-QDPLWQTTMPPQAF | 370 |
| *Acinetobacter_baumannii*_d | KRNITFALGTECLTHQDAY-VSSILSFVLIRKYA----SNLYQLD-HGEF---------- | 416 |
| *Acinetobacter_baumannii*_118362 | GGSITFAMGTEFFTHVDAQ-LGGIIQDSISRTYV----SNLYQMD-DAIF---------- | 404 |
| *Acinetobacter_baumannii*_e | GGSITFAMGTEFFTHVDAQ-LGGIIQDSISRTYV----SNLYQMD-DAIF---------- | 405 |
| *Acinetobacter*_sp._a | KRSITFAMGTEFLNHTDAQ-IHPLLIEQFSRTYV----SNLYQYD-QSIF---------- | 392 |
| *Acinetobacter_guillouiae*_NIPH_991 | SGSISFAMGTEFLTHDDAL-IHPLMVEPLARSYI----SNLYQYD-QSIF---------- | 392 |
| *Trinickia_fusca* | PRSVGFALGDERIEHEDFV-LEGPLPIVWQRTYR----SFFDANDAHGEL---------- | 416 |
| *Burkholderia_contaminans* | IGSISYSFGDETFSHADFE-LPGAMPVVWMRVYR----SRLAGYD-DGEL---------- | 374 |
| *Burkholderia*_sp._Bp9004_c | TGSIDFAFGDETFTHTDFE-LPGALPLVWARTYR----SRLSAYD-TREL---------- | 398 |
| *Burkholderia*_sp._Bp9142 | KASIDFAFGDETFTHTDFD-LPGALPLVWARTYR----SRLSAYD-NGEL---------- | 398 |
| *Burkholderia_cenocepacia* | EASIDFAFGDETFSHVDFD-LPGAMPLVWERTYR----SRLSAYD-NGEL---------- | 399 |
| *Burkholderia*_sp._4701 | ------------------------------------------------------------ | 330 |
| *Burkholderia*_sp._AU6039 | IGSISFARGAERIDH--------------------------------------------- | 374 |
| *Burkholderia_stagnalis* | GRAIGLALGDESFTHTDFA-LPGVVPVEWARTYR----SNFGAHDEQGPL---------- | 407 |
| *Burkholderia_reimsis* | GGAIGLALGDESFTHTDFV-IPGVMP---------------------------------- | 386 |
| *Paraburkholderia_rhizosphaerae* | TGSIGFALGDESFTHTDFV-LPGVLAVEWVRAYR----SNFDAHDTAGPL---------- | 405 |
| *Pseudomonas_putida* | NCSISFATGSETIVHTDFQ-LPGAFPIEWYRTYR----STLSAFD-DSPY---------- | 367 |
| *Burkholderia*_sp._Bp9004_d | ------------------------------------------------------------ | 314 |
| *Burkholderia*_sp._Bp9004_b | GS---------------------------------------------------------- | 317 |
| *Burkholderia*_sp._Bp9004_a | GSNISFATGSESLTHTDFI-LPGPFPIAWTRSYR----SSQAAYD-SGEL---------- | 389 |
| *Burkholderia_cepacia* | GGSISFATGTETLSHTDFV-LPGPFPIVWTRTYR----SSLGAYD-AGEL---------- | 390 |
| *Pseudomonas*_sp._LP_8_YM | GSNISFATGSESFTHTDFS-LPGCFPIEWARTYR----SSLSALD-EGPL---------- | 361 |
| *Pseudomonas*_sp._FSL | GCSISFATGSETVAHADFS-LPGPFPVEWVRTYR----SSLDALD-HGSL---------- | 362 |
| *Pseudomonas*_sp._URIL14HWK12:I1 | CNRISFARGSEQITHTDFT-LPGPFPVTWTRNYR----SSLSALD-TGPL---------- | 375 |
| *Pseudomonas*_sp._FSL_R10-0765 | GHSISFARGSEYLAHTDFT-LAGPFPIEWSRAYR----SSQSALD-SGAF---------- | 371 |
| *Pseudomonas*_sp._c | GHSISFARGSEYLAHTDFT-LAGPFPIEWSRAYR----SSQSALD-NGVF---------- | 371 |
| *Pseudomonas_helleri* | GHSISFARGSEYLAHTDFT-LAGPFPIEWSRAYR----SSQSALD-SGVF---------- | 371 |
| *Pseudomonas_helleri*_b | GHSISFARGSEYLAHTDFT-LAGPFPIEWSRAYR----SSQSALD-SGVF---------- | 371 |
| *Andreprevotia_lacus* | KNSIDFAMGTESVSHVDFV-LNGVFPLQWSRTYR----SNLAAFN-DSEQ---------- | 398 |
| *Pseudomonas_lundensis* | CNSILFSLGAETFSHTDFS-LPGPFPIDWTRT---------------------------- | 367 |
| *Pseudomonas_graminis* | IKSISFAMGSESVSHTDFS-LPGPFPIEWTRTYC----SSLDAYD-QDII---------- | 406 |
| *Pseudomonas*_sp._a | IGSISFAMGSESVSHTDFS-LPGPFPIEWTRTYC----SSLDAFD-EDVV---------- | 400 |
| *Pseudomonas*_sp._b | GNSISFAMGSESVRHTDFS-LPGPFPIEWARTYC----SSLDAYD-QDVI---------- | 400 |
| *Pseudomonas_cichorii* | CNSISFALGSETFSHTDFS-LPGPFPIHWARLYN----SRLSAYD-QGIM---------- | 394 |
| *Pseudomonas_sp._WS_5354* | ------------------------------------------------------------ | *0* |
| *Pseudomonas_helmanticensis* | ------------------------------------------------------------ | *0* |
| *Neisseria*_sp._HMSC065C04 | GRAVAVQKTKGAVEQISNK-ITGLIGEHMADYWMLEQVGGKARHDHGGTA---------- | 368 |
| *Xanthomonas_sacchari* | PLHAAVQVTRKAYQNIAKR-QKGLIGEHMADYHELKRLGGHWPHDNAKGHWS-------- | 328 |

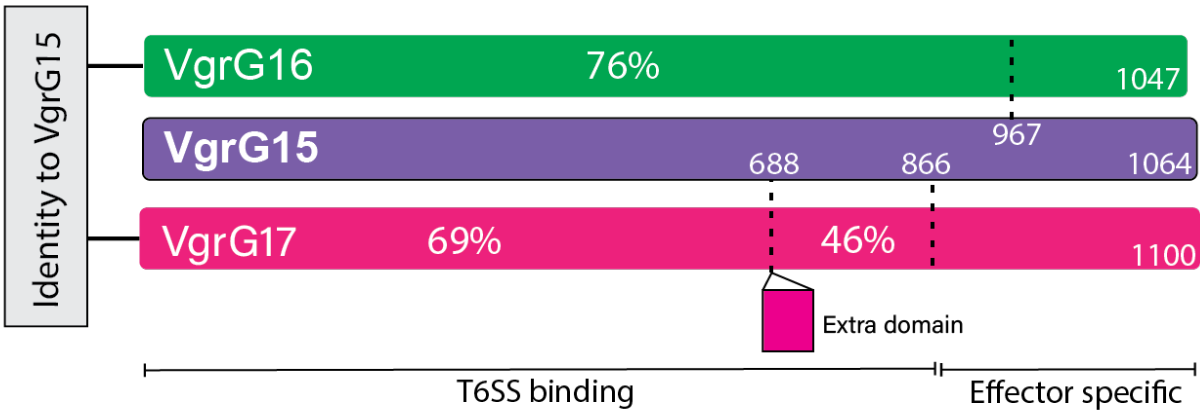


**Supp Figure 12. The VgrG15 N-terminal domain is highly conserved as compared to the N-terminal domains of the other *A. baumannii* AB307-0294 VgrG proteins.** Green indicates VgrG16, purple VgrG15 and pink VgrG17. The amino acid length of each is indicated by the number at the end of each schematic. Percentages show sequence identity between either VgrG16 or VgrG17 and VgrG15 in the regions between where the dashed lines are placed. The small pink box below VgrG17 denotes an extra domain present in VgrG17 that is not found within the other VgrGs. Overall, we predict that amino acids required for overall T6SS structure are within the N-terminal region, while the C-terminal region is specific for interaction with the effectors.

### Tse15

MAKENTQQKEIKRQPAVELAVFNLNSVTDVADLQMIASQVQLYLQVCGNTTLEQIKSKANITTVANIFALTGSVLDLMLYATDKKTGDAAVQR
**GALLAANLIGLFSEPNNEAHARMAIRPMFGLMAECLYR**ENGKIKETDIKR**LGLHLNAMIAGDLENFLK**ETQAKLSSLLISATTLGVTILQSMA
TPATGINAGITTAAGASAEKRDPKLK**FTNWAVPLIDLLGKPSQANLTPKIQPNITSRLQQEATQAIAALSQTLQQQANAGQKYTIAWLLQETL**
**KAIQALENKGNASVPINQTGEYER**HTK**GDTLEFVSLQADALNAPPCEGADSQSGK**SISYSIGAERVQHADFYLPKIGFSFIRQYNSQMDEFDQ
SMVGARWMMPFSNMIQQNAQGYLFIDSKGRKHQLPVSIIFETYEVPYEGWIIKPLKNGELILDFGGEWRSHFQSFDGGKNYYLVKKMNETSQE
EILLEYLLLDHIAYLKVINFKLKQAEYELKFAFNEQVKIIAVFLDDKAEPLAR**YEYDTQGNLIK**AIDQNGHTRTYEYNQFHQLTRYTDRTGRG
QNIRYESTEAKAKAIEEWADDGSFHTKLKWHPRLRQVAVYDAYDVPTYYYFDLDGFTYRTRLADGRESWYSRDGKKRITR**QIDFDGR**ETQQEY
NDQDQLVKIVQPNGGIIRFAYNKQGNLVEIKDPEGSIWKREYDENRNVSKEINPLGHITQYKYNNDNQLVEVIDAKGGVKKIQYNELGQMISY
TDCSGKSSTWEYDEDGALTAEQTANNKVVQYFYSTKGRDKGQLQSIIYPDGLKEYFEHDEEGRLLKHTDTKGLVTEYKYNQVGLLEQRIDANR
HSVAYQWDKQGRIQKLINQNQAEYLFGYNPYGYLIREQAFDGEEKHYSYNENGRLFQIRRPNILTQFDYYADGQIASKSFTHLHTGQKQTEQF
DYNLNSQLSRASNEVSQIDLYRNALGQLVREHQHYKIPELKPLTAVLHYEYDELGNLIKTIRPDGHTLNHLVYGSGHIYAIGLNNQEVVSFQR
DDLHRETTRLLANGLMQTKQYNDVGLLSSQFIQPEQETQDYLQYQAHRKYHYDKNYLLSQVEDSRLGKLNYQYDPIGRLIAAQSLHKTESFNF
DPAGNLIDSESVLSPAQIKNNLIKSYKGKHYQYDVQGNVTEIIQAGKNLKLTWDNQNRLIRSDNNGLVTEYGYDVFGRRLYKKTAKELTLFGW
DGDLMIWESFKSAQTNYTKHYIYEPDSFVPLLQAGYKDFIQLIETPDYQEYQTKPYSIYKDPVWNRNLGKERTALEQFTFYHCDQVGTPQTMT
NIRGECVWEILQDTWGAVSQIKALNQDNPFEQNNLRFQGQYYDRETELHYNRYRYYEPHSARYVSKDPIGLEGGMNTSSYVSDPNQWIDPKGL
NSFNYGEMFGIPASAQSGLAYQGQR**NYECYAETGELCK**IKVPPLFDYVACSGGGLGIGVGFVK**NQWTGEYYISGSKDSLLIPVAK**SVAQNKQF
SAK**DLAGASCVGGNIHNIPSYTK**TTMTMGEITNEFVSGASVTVGGGAYGAVANVVVPLVSKSSPVKGTWASELGVGTPGFNVGVSGTVSVDTI
LDAVKPSKK

### VgrG15

MFNNIFQILESFGFLSQHR**SVYLQFSDASLNSQVFLQRIDGQHYLNQGMTAELICLSTNAHIPLKTFIGLQVAVDQVTDR**GSFFR**TTGIITGA**
**SQGQSDGALTLYKLTVSDPTYLWHK**RRNSRVFMNKSVK**EISEILFQEWQGKSPLFASSLTLDLSGLKQTYDVRPFVMQLNESDYDFLTR**LWR**S**
**EGISWLIDEAELTVASNMDNIQPQK**LR**LIDDNNQYQALTR**RVIRYHR**SSATEQFDSMTSLMADRSLQPTSIFVQRWQPDVLQQTDGAGSVQSK**
**HQHSTNYDNQSLSLEEAWHFSPAWMQDLNGEDGATSASNQQIEKFNQNLSAYYDAQSK**QFIAKTTVR**DTQVGYWFELNEHPEIDQHESTDKEF**
**LIIGKNYYNQNNLPKDLNQQIQTLLQQSDWQASNTDERQANQLILQR**R**YIPTTPAYNPQTHSPVAHPQR**AK**VVGPEGEEIYVDEWGR**IKVRFL
FTR**SDDHSHDGGAGTNNNDTDSAWIDVLTPWAGEGYGAR**FLPRIGEIVVIDFFNGDIDRPFVMGRIHEAQRHPTKFDNKGKLPDTKKLSGIRS
K**EVSGSGFGQLRFDDTPGQISTQLQSSHGASQLNLGK**LSHPK**DKAESEDRGEGFELRTDQWGALRAGQGLLVSTHK**QDNAK**GDHLDAEVAK**KQ
LEGSQTNSKALSDIAK**NQKTDEIESIEQLKDFASQIQQQIAK**FEKALLLLSSPDGIALSSSEDIHISADAQINQIAGDSINISTQK**NVIAHAQ**
**NRLSLFAAQSGLK**AVAAQGK**VEIQAQADALDVLSKLGITISSTDDKVIISSPK**EVK**ITGGSSQITLNGSGIFPK**TGGK**FQVNAGQHLFMGGAS**
**ANASAPELPKAKPMQGALELLRSYGGDNFFK**QNSYKVIDSLGKQITGK**LDGNGFAQVTGIAPGPAK**VVFEK**DNTSAWLQSSDFKRNYTWAEPV**
**K**SVQGLMK**NALEAVGQNTMSQLQNNLLSTDKNSFK**NLGK**NTLDNLAGQTVAQIKNQVTNTALNTVSKQLNLNLSADQMK**SLGQMATNPSQSLE
MLK**EQGGDFLSDQMTAK**LFK**TTNQESPIQQGDLDTFVR**SKK

### Tde16

M**SQNSVVAPLNTFSPK**DLTAKKAEVDKWFR**EYTNGVVTVDR**LETICRSVPVLGSAFAIGDIIIDIISMINKGGLDKVEIFDWLNLGIDVIGLV
PMGPVGPSVRSAARPALFYVKNESEKIIKAQAKKLGKKTLTSQEVKKALSTGFKDSASVFLTTIIAENVAGTLENFAKKGQSLLNQILKEVGN
WIVLLTKTIDDGFKKLVSGSLNGLPNLKRAGQQSLGVIK**GIFELDGTR**IVNNAKYATENVAKTVGK**GYVNLANLAVSDEAR**AKVLALGAKIRS
IGQVAQAKVNGLSDPNTLWTIGWLFSIMGMVAAKHRQKRAQIKAKETTKANASHPSTATDKSNKQAHAENNANQCKNCMGGTGGSITFAMGTE
FFTHVDAQLGGIIQDSISRTYVSNLYQMDDAIFGARWVTPFTTKISRKFKYTSKKKDHKDYLNGLEYICLDGRAIDLPDLKKGQSIYDPIEQY
TYTVLSDQLHLIAYGEDEKRYYEKYGEDYRLSYIERKNGFKVALRYDHVSTDNKTILSDILFKQDDNLLAHLALQLTPQGLVSDIWTIKNGQL
DRVLASYDYDQQGDLVQATNEFAASYYYQYTHHLITRYTDLTHRGMNLKWDGILPTSKAIEEWADNASRASKLEWDKNIRKTTVLDVEGNSTE
HYYDIDGYTYRIVYPDNFEEWFFRDDAKNITLHIAKDGSKTSYTYDERGNVLTTTQDDGATSYFEYDEKNQLTGMVDAEQGRWFKQYDGSGNL
IKEIDPLKHETAYVYNAMGLVTSITDAKGGSKSLKYDDQGNLISYTDCSGKETKWQYDERGRVISIENALNQKVEYFYTELTLENREPIIKGL
PLNAFGQLEKIKHADGTEEHFIHDAEGRLLAHVDPKQNITRYEYDEAGLILSRTDALNHKLKYKWDRLGRLTRLINENGASYQFFYDVASRLV
KEIDFDGKETVYHYDEKSGQLATSIEVASAYGQDLKDRAAPKDRIQQFIFDSMGRLEQRTAGYGHYGLELEEKQTEEFAYDYMGRIIQAKNAQ
SNLQWFYDAAGNLVQEHQQDYKINKTAVWKHQYDEINDRIKTTRPDGQVIDWLTYGSGHVQSLIVNGQDFVSFERDDLHREIARHYANGVSQE
QQYDLAGRLKSQMMLSEHENGYQNQYKRHNNALEQTSQLVQRLYQYDKTGELTAIRDTRRGNIAYKYDPVGRLLEASSKLGKETFSFDPASNI
LDSYHSQKVQSHSQKLDETSYGYNRLVNNVVKEYLDQQYQYDAYGQLIRQKTSQGDLNLEWDVYGRMVKSRNSQYTAEYRYDALGRRIQKWSK
HHHTGQEQNIIYGWDGDTLAYESTEELTKHYIYEKDSFVPMLQAVYLSPIELHQTPDWSDRPYNIHRDPLWKTEKEGKEFDDVWFYHCDHLGT
PQEMTDHTGAIIWKAEYKAWGECKAEEAKSNFFENSEIISNNIRFQGQYFDEETGLHYNRYRYYSPYVGRFVSKDPIGLLGGNNVYVYAKNPI
TWIDSKGLCSTTLNRNLGGVKGDHLQAHHIIPEEIWAK**RKDFLDDIGIGGNRDKAENGVLMPDSEAK**AKQMKR**QLYHCGSHPIYSAGINQK**LG
QIQREFESKKITASQAR**DKVANLQSSMRLVLITPGTKPIR**LS

### VgrG16

**MLNSIHQVLDSLGISPQKR**AIHVQFTSSILNAQVFLQR**IDGVHALNDGLK**AELLCLSTNATIQLK**SFIGVQAAVDIVTER**GELTR**VTGIITHA**
**QQGQSDGSLTLYKLTLEDPTALWK**YRRNSRVFMNK**SVVEIWEILFK**EWQTKNPLFAASLSLDLSGLTQTYDVR**PFVMQHNESDWNFLTR**LLR**S**
**ENISWLIDEAQHIVPSTETPIQAQK**LR**LIDANSQYQPLDRK**TIRYHR**SSAVEQYDSMTR**LTAER**SLQPNVMHIQRWQAEILDQEEGIGSVQSK**
**HQHSEHYDNATLGLEQAWNYSPAWIGDLK**GEDGVTKSGNQQVER**LNQNLNNYYEAQAKRFIAQTTVRDAYVGYYFELNEHPEIDQHESTDRSF**
**LIISKSFFNQNNLPK**DLNDR**INGLLAQSNWAIQNPENSDER**QANQLILQRRHIPTTPAYSPQIHSPVTHPQRAKVVGPEGEEIYVDEWGRIKV
RFLFTRSDDHSHDGGAGTNNNDTDSAWIDVLTPWAGEGYGVRFLPRIGEIVVIDFFNGDIDRPFVMGRIHEAQRQPTKFDNKGKLPDTKKLSG
IRSK**EVSGGGFGQLRFDDTPGQISTQLQSSHGASQLNMGK**LSHPKDKAESEDRGEGFELRTDQWGALRAGQGLLVSTHKQDNAK**GEHLDAEVA**
**K**KQLEGSQTNSKALSDIAKNQKTDEIESIEQLKDFASQIQQQIAKFEKALLLLSSPDGIALSSSEDIHISADAQINQIAGDSINISTQKNVIA
HAQNRLSLFAAQSGLKAVAAQGK**VEIQAQSDALDVLANKGITISSTEDCIEISSPK**EIVITGASSQITLNGSGIFPKTGGK**FQVNAGQHVFQG**
**GASASVKSSLPPPPK**R**AQGVLELLHDYSHGAFVKGAGYTVTDNLGK**VVNGKLDDKGFAR**VSGLATGSVKVLFDPDPRNPWDEASDFKRKVEWP**
**NQNDVEGSIGIESGSK**GLTETLKNDFK**KLSQLASNVK**SVQQKIESVKKIKDQGAK**ALLPMAMEQVLGGSDGSKLSQSFVSQNSFMPSSQQSNP**
**LRR**LAQASSDKTAITK**IQSPFETFV**

**Supp Figure 13. Peptide signatures of Rhs effectors and their cognate VgrG within the secretome of T6SS active *A. baumannii.*** Peptides identified by MS/MS are in bold and underlined. The top shows coverage for Tse15 where the domains are coloured: N-terminal clade domain is orange, Rhs domain is grey and the toxin domain is blue. The second sequence is VgrG15 where the entire sequence is purple. These colour schemes are retained for Tde16 and VgrG16 where the Rhs protein (Tde16) is coloured into the three domains and VgrG16 is purple.
